# Genetic approaches in mice demonstrate that neuro-mesodermal progenitors express *T/Brachyury* but not *Sox2*

**DOI:** 10.1101/503854

**Authors:** Dorothee Mugele, Dale A. Moulding, Dawn Savery, Matteo A. Molè, Nicholas D. E. Greene, Juan Pedro Martinez-Barbera, Andrew J. Copp

## Abstract

Neural tube and somites have long been thought to derive from separate germ layers: the ectoderm and mesoderm. This concept was challenged by the discovery of neuro-mesodermal progenitors, a bi-potent cell population that gives rise to both spinal neural tube and somites. In line with their proposed potency, these cells are considered to co-express the neural marker *Sox2* and the mesodermal marker *T/Brachyury*. We performed genetic lineage tracing in mouse embryos and confirmed that *T*-expressing cells give rise to both neural tube and mesoderm. Surprisingly, however, Sox2-expressing cell derivatives colonise only the neural tube after embryonic day 8.5. Deletion of *Sox2* in *T*-expressing cells was compatible with an otherwise normal neural tube and paraxial mesoderm. Moreover, *Sox2* expression is absent from the chordoneural hinge, where neuro-mesodermal progenitors are located. Our findings demonstrate that neuro-mesodermal progenitors express *T* but not *Sox2*, suggesting the need for re-evaluation of the neuro-mesodermal progenitor hypothesis.

## INTRODUCTION

The concept of neuro-mesodermal progenitors (NMPs), as a bi-potent progenitor cell population that gives rise to neural tube and paraxial mesoderm, has attracted a great deal of attention in recent years. While neural and mesodermal lineages were long assumed to segregate during gastrulation, an increasing number of studies suggest that NMPs exist and function during post-gastrulation development in mouse^1–3^, *Xenopus*^4, 5^, axolotl^6^, avians^7^, zebrafish^8^, and humans^9^.

The lack of cell type-specific markers makes it difficult to study NMPs. Yet, the discovery of cells expressing both *T/Brachyury* (here referred to as *T*) and *Sox2* in the region where NMPs are located has led many researchers to conclude that these are the NMPs^6, 9–11^ Indeed, co-expression of master regulators of mesoderm (*T*) and neural (*Sox2*) development fits well with the proposed potency of the dual-fated progenitors. Based on this assumption, *T/Sox2* double-positive cells have been studied extensively *in vitro*^12–15^ and *in vivo*^6, 9, 11, 16–20^. Despite the increasing interest in NMPs, their precise role in embryonic development still remains to be elucidated, though it is hypothesized that they are essential for body axis elongation^2, 9, 12, 16, 20–22^.

The literature is often inconsistent regarding the embryonic location of NMPs and the associated nomenclature. For this study, we adopted the definitions provided by Cambray and Wilson^1, 2^, who performed grafting experiments in mouse embryos. According to their work, a cell population which gives rise to neural tube and paraxial mesoderm is located at the border between node and primitive streak remnant, the so-called node-streak border (NSB), from embryonic day (E) 8.5 onwards (Figure 1a). In addition, these authors found cells with similar characteristics when homotopically grafting tissues spanning regions 1 – 3 of the epiblast that bilaterally flanks the primitive streak remnant, termed the caudo-lateral epiblast (CLE). After internalisation of the node around E9.0, NMPs can be found in the chordo-neural hinge (CNH), which lies in the midline of the tail-bud, directly caudal to the forming hindgut and notochord (Figure 1b).

**Figure 1.**
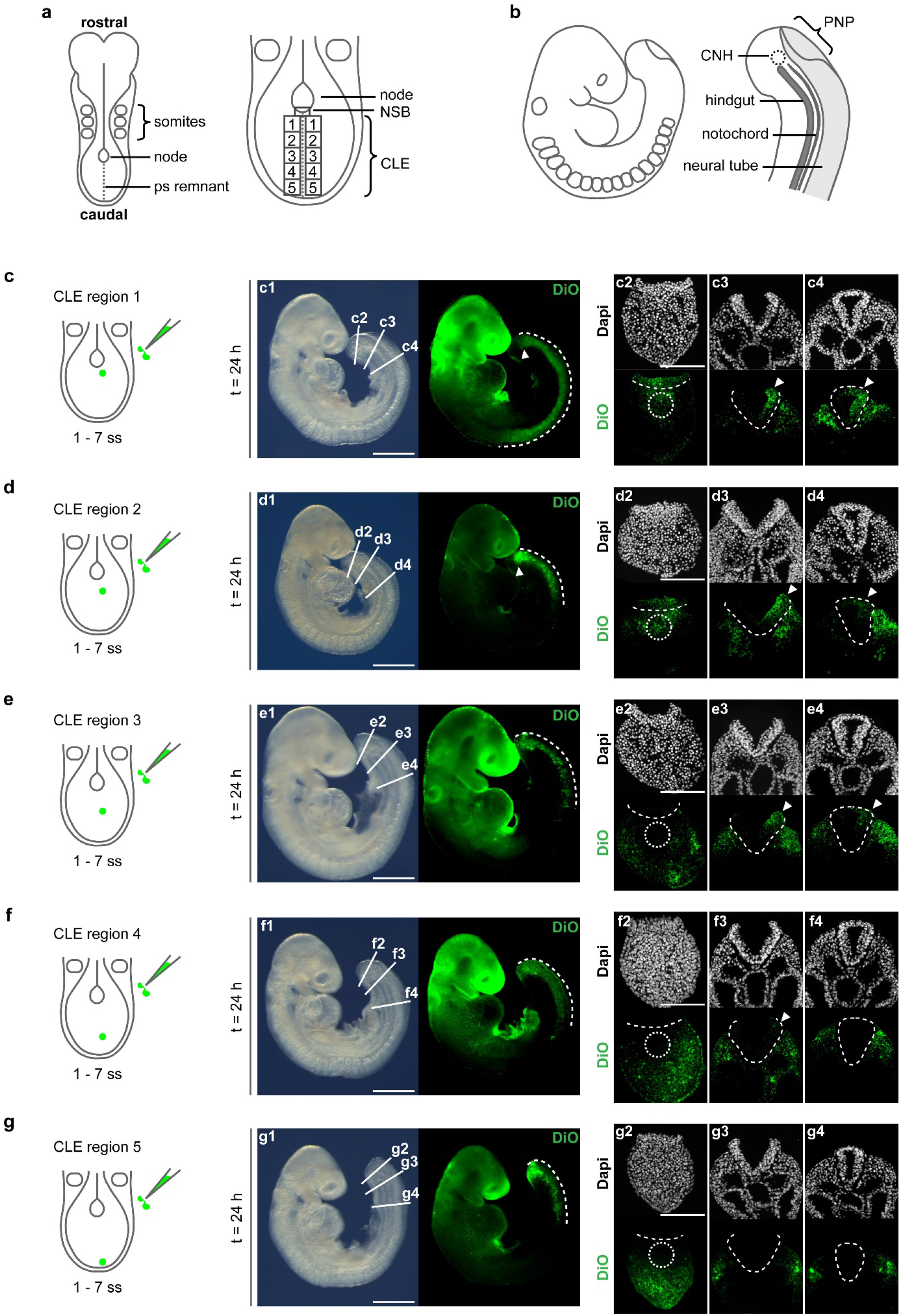
NMPs are located in the rostral CLE and colonise the dorso-lateral neural tube. (**a, b**) Schematic representations of E8.5 and E9.5 mouse embryos. At E8.5 (**a**), longterm progenitors for the neural tube and paraxial mesoderm reside in the NSB, which is located at the rostral end of the primitive streak (ps) remnant, directly caudal to the node. The CLE is located caudal to the NSB, flanking the primitive streak remnant bilaterally. At E9.5 (**b**), NMPs are located in the CNH, directly caudal to the elongating hindgut and notochord, and underlying the posterior neuropore (PNP). (**c – g**) Cells in regions 1 – 5 of the CLE were labelled with DiO in E8.5 embryos followed by 24 h whole-embryo culture. DiO-positive cells are specifically retained in the CNH after labelling regions 1 and 2 (white arrowheads in **c1, d1**), as confirmed by sections (dotted circles in **c2 – g2** indicate CNH). White dashed lines in **c1 – g1** indicate how far rostro-caudally labelled cells contribute to axial tissues. Sections (**c3 – g3**; **c4 – g4**) show that labelled cells colonise the dorsal-lateral neural tube (white arrowheads), not the ventral and ventro-lateral regions. White dashed lines in **c2 – g4** outline the neuroepithelium. Contribution of labelled cells to neural tube decreases and contribution to mesoderm increases with more caudal DiO labelling (**c2 – g4**). See Supplementary Table 1 for summary of DiO-labelled cell colonisation patterns. Number of embryos for labelling of regions 1 – 5 respectively: 8, 8, 7, 7, 8. Scale bars: 500 μm for whole embryos (**c1 – g1**); 100 μm for sections (**c2 – g2**).

The initial aim of this study was to define the role of NMPs in neural tube formation, since it is not clear to what extent the progenitors contribute to this tissue. We traced cells either from the NSB or CLE into the closing neural tube, and found that the caudal end of the embryo harbours two distinct populations, both of which fulfil the criteria of NMPs, yet give rise to different domains within the closed neural tube. NMP function was addressed using laser ablation and a genetic approach, based on the proposed co-expression of *T* and *Sox2*. These experiments reveal that cells expressing *Sox2* after gastrulation give rise to neural tube only. Therefore, we propose a new model of T-positive NMPs, which up-regulate *Sox2* only as they adopt a neural fate.

## RESULTS

### NMPs are located in the rostral CLE and specifically give rise to the dorsal neural tube

Cells were traced from regions 1 – 5 of the CLE, to better define the location of NMPs in E8.5 mouse embryos. A small number of cells were labelled by injecting the green fluorescent dye DiO into the region of interest, followed by whole embryo culture for 24 h (Figure 1c – g). Embryos with 1 – 7 somite pairs (E8.5) were used since at this stage the caudal border of the node is clearly visible, enabling reproducible injections. Release of DiO into the amniotic cavity, which occurs during injection, caused non-specific labelling of the head-folds but not of the caudal region where cell tracing was conducted (Supplementary Figure 1). Labelling of CLE region 1 produced a colonisation pattern as expected of NMPs: DiO-positive cells were present in both neural tube and somites, over a considerable length of the body axis, as well as being specifically retained within the CNH (Figure 1c). Labelling of region 2 also resulted in accumulation of DiO-positive cells in the CNH, although a shorter length of the body axis was typically colonised (Figure 1d). In contrast, DiO injection into regions 3 – 5 did not lead to CNH colonisation: only neural tube and paraxial mesoderm contained DiO-positive cells and only over a short axial length. As described previously^2^, the contribution of DiO-labelled cells shifted towards mesoderm, and away from neural tube, as more caudal regions of the CLE were labelled, so that injection of region 5 yielded mesoderm labelling only (Figure 1e – g; Supplementary Table 1). Notably, when the neural tube was colonised, labelled cells were always located in the dorsal and dorso-lateral domains, never in the ventral neural tube (white arrowheads in Figure 1c – f). This was independent of the position where dye was initially injected in the CLE (regions 1 – 4).

After labelling region 1, DiO-positive cells translocated in a ventral-to-dorsal direction and reached the dorsal neural tube after 6 – 9 h (Supplementary Figure 2). Even after prolonged culture up to 45 h, DiO-labelled cells were detected in the dorso-lateral domain of the closed neural tube only (data not shown). In contrast, when DiO was injected directly into the lateral portion of the closed neural tube of E9.5 embryos, the dye-labelled cells spread to some extent, but did not reach the dorsal domain (Supplementary Figure 2). Hence, the ventral-to-dorsal translocation in the neural tube is specific to cells arising in the CLE.

**Figure 2.**
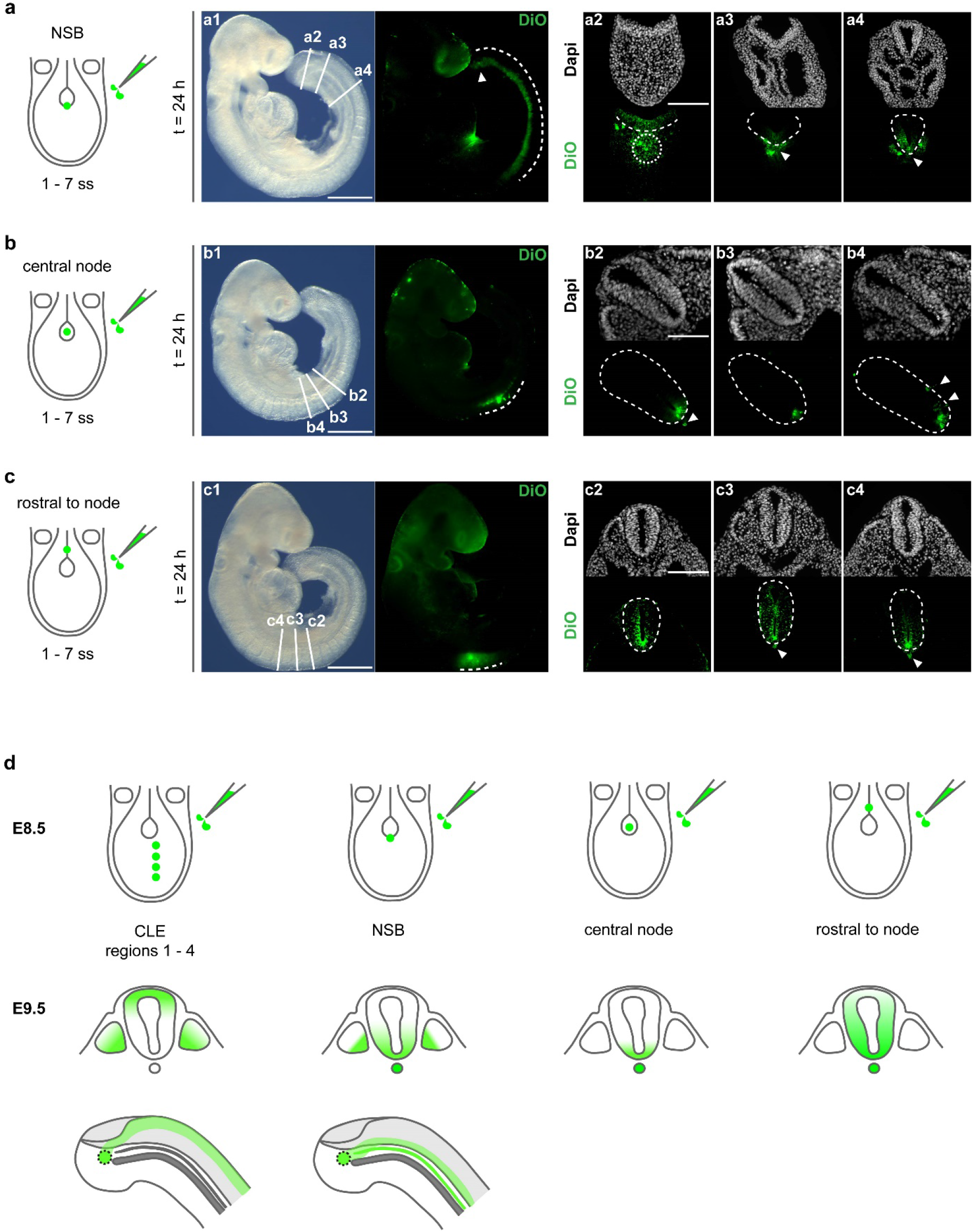
Ventral and ventro-lateral domains of the neural tube are derived from cells located in and around the node. DiO-labelling of cells in the NSB (**a**), the centre of the node (**b**), and rostral to the node (**c**) in E8.5 embryos followed by 24 h whole-embryo culture. White dashed lines in a1 – c1 indicate how far rostro-caudally labelled cells contribute to axial tissues. (**a**) NSB injection yields DiO-labelled cells that are specifically retained in the CNH (white arrowhead in **a1**), as confirmed by transverse sections (CNH: white dotted circle in **a2**), the notochord (white arrowheads in **a3 – 4**), paraxial mesoderm, and the ventral neural tube (**a4**) in 9/9 embryos. (**b**) Cells labelled in the central node colonise notochord (white arrowhead in **b2**), floor-plate (**b2 – 4**), and a few cells in the lateral neural tube (white arrowheads in **b4**) in 8/8 embryos. (**c**) Cells labelled rostral to the node colonise notochord (white arrowheads in **c3 – 4**) and the neural tube along its dorso-ventral axis (**c2 – 3**) in 8/8 embryos. (**d**) Summary of DiO-labelling experiments. The bottom row shows a schematic representation of the caudal region in lateral view indicating the variation in neural tube colonisation by cells labelled in CLE region 1 (left) and NSB (right). See Supplementary Table 1 for summary of labelling patterns. White dashed lines in **a2 – c4** outline the neuroepithelium. Scale bars: 500 μm for whole embryos (**a1 – c1**); 100 μm for sections (**a2 – c2**).

### The ventral neural tube is derived from cells located in and around the node

As CLE injection labelled only dorso-lateral neural tube, we next investigated the origin of the ventral neural tube, including the floor-plate. Cells were labelled at more rostral positions: in the NSB, the centre of the node, or directly rostral to the node, followed by embryo culture for 24 h. NSB labelling led to colonisation of notochord, floor-plate, ventro-lateral neural tube, and sclerotomes along considerable lengths of body axis (Figure 2a). Labelling of the central node (Figure 2b) contributed DiO-positive cells to notochord and floor-plate, consistent with other studies^23, 24^, but only along a short axial length. In addition, some contribution was observed to the ventro-lateral neural tube. Tracing cells from directly rostral to the node (Figure 2c) revealed colonisation of notochord and neural tube along its entire dorso-ventral axis. However, DiO-positive cells contributed only to a relatively short length of the body axis. While dye-labelled cells were retained in the CNH after NSB injection, this was not observed after labelling the central node or rostral to the node (Figure 2a – c; Supplementary Table 1).

Hence, we observe NMP-like cell behaviour (retention of labelled cells in the CNH and colonisation of neural tube and mesoderm along a considerable axial length) after labelling both NSB and CLE region 1 (Figure 2d). However, the NSB gives rise to ventral neural tube and notochord, whereas region 1 cells colonise the dorso-lateral neural tube but not notochord. As CLE region 1 most closely resembles the NMP colonisation pattern described in the literature^10, 12–16, 20^, we focussed on this region in subsequent analyses.

### Ablation of the rostral CLE transiently affects neural tube and somite formation

To determine the requirement for NMPs in early embryonic development, we performed laser ablation of the CLE region 1 in E8.5 embryos (Figure 3a). Following 24 h culture, ablated embryos showed a trend towards smaller somite number and body length compared with non-ablated controls. However, the axial length of open spinal neural folds (the posterior neuropore; PNP) was significantly larger, indicating a delay in neural tube closure, although there was still considerable overlap between ablated and non-ablated embryos (Figure 3b – d). Serial histological sections revealed a normal neural tube along most of the body axis of ablated embryos (Figure 3e2, 4), but a short stretch of ~ 3 – 4 sections per embryo (~30 – 40 μm) around forelimb bud level showed a highly malformed neural tube (Figure 3e3). Whole-mount *in situ* hybridisation (WISH) against the sclerotome marker *Pax1* did not reveal any obvious effect on somite formation, apart from a missing somite in 1/8 embryos (Figure 3f5). *Dll1* expression in pre-somitic mesoderm was similar to controls after 24 h culture (Figure 3g1 – 4), although expression was clearly reduced in the tail-bud region of embryos cultured for only 6 h following laser ablation (Figure 3g5 – 8). Hence, ablation of CLE region 1 transiently affects formation of both neural tube and paraxial mesoderm.

**Figure 3.**
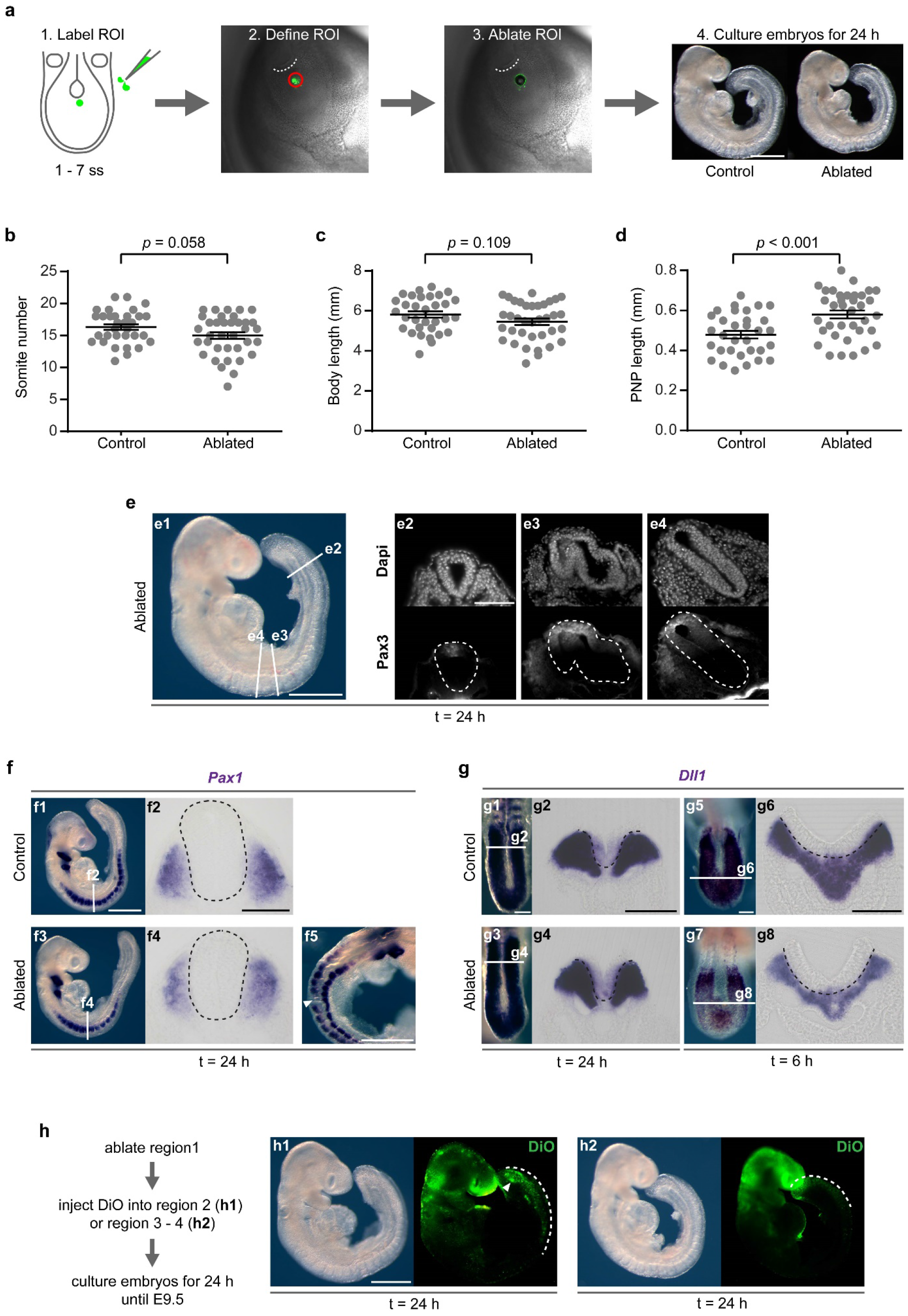
Laser ablation of CLE region 1 transiently affects formation of neural tube and somites. (**a**) Summary of procedure for laser-ablating region 1 of the CLE. White dotted lines: caudal border of the node; ROI: region of interest. (**b – d**) Embryo measurements after 24 h culture (n = 32 control, n = 36 laser-ablated). Data shown as mean ± SEM including the individual data points. Unpaired, two-tailed Student’s *t*-test. Laser-ablated embryos do not differ significantly from non-ablated controls in somite number (**b**), or body length (**c**), but have longer PNPs (**d**). (**e**) Although the neural tube is unaffected over most of its length (**e2, e4**), a short stretch of ~30 – 40 μm has severely abnormal morphology (**e3**, n = 13/17 embryos). The white dashed lines (**e2 – 4**) outline the neuroepithelium. (**f**) WISH for the sclerotome marker *Pax1* shows no marked difference in expression between control (n = 10) and laser ablated embryos (n = 7/8). One laser-ablated embryo, however, had a somite missing (white arrowhead in f5). (**g**) WISH for the presomitic mesoderm marker *Dll1* reveals no difference in expression between control (n = 7) and laser-ablated embryos (n = 11) after 24 h (**g1 – 4**). Yet, embryos cultured for 6 h only after ablation of region 1 (**g5 – g8**) show strongly reduced *Dll1* expression (n = 10/10) compared to controls (n = 9). Black dashed lines in **f2, f4, g2, g4, g6** and **g8** outline the neuroepithelium. (**h**) After ablation of CLE region 1, the CNH is re-populated following 24 h culture by cells labelled in region 2 (white arrowhead in **h1**, n = 7/8 embryos), but not by cells labelled in regions 3 – 4 (**h2**, n = 0/8 embryos). White dashed lines in **h1 – h2** indicate how far rostro-caudally labelled cells contribute to axial tissue. Scale bars: 500 μm for whole embryos (**a, e1, f1, f5, h1**); 100 μm for caudal regions (**g1, g5**) and sections (**e2, f2, g2, g6**).

The short-lived effects of CLE ablation suggested embryos are able to compensate for the loss. To address this possibility, DiO was injected either into region 2 or regions 3 – 4 of the CLE directly following ablation and before 24 h culture (Figure 3h). Cells from region 2 colonised the CNH in ablated embryos (white arrowhead in Figure 3h1), but this was not observed after labelling regions 3 – 4 (Figure 3h2). In addition, a longer stretch of the body axis was labelled than observed after DiO injection into region 2 in non-ablated embryos (Figure 1d1). This suggests that region 2 cells re-populate the deleted region, and thereby attenuate the effects of ablation. Having demonstrated rapid recovery following ablation of the rostral CLE region in which NMPs are proposed to reside, we used genetic lineage tracing to further investigate fate and function of *Sox2-* and T-positive cells.

### Mesodermal contribution of *Sox2*-expressing cells from E8.5 is a rare event

If NMPs co-express *Sox2* and T, the descendants of either lineage should colonise both neural tube and somites when traced from E8.5 onwards. To test this prediction, a *T^CreERT2/+^* driver line was crossed with the *Rosa26^mTmG/mTmG^* reporter, with activation of CreERT2 recombinase by tamoxifen injection at E8.5, and embryos harvested from pregnant dams at E9.5 (Figure 4a). As observed previously^25^, GFP-positive cells were abundant in neural tube, somites, notochord, and CNH, consistent with NMPs being 7-positive. In line with previous reports, an estimated 85 – 90% of all cells in the neural tube were GFP-positive^25^. Interestingly, GFP-expressing cells were also present in the hindgut (white arrowheads in Figure 4a3 – 4).

**Figure 4.**
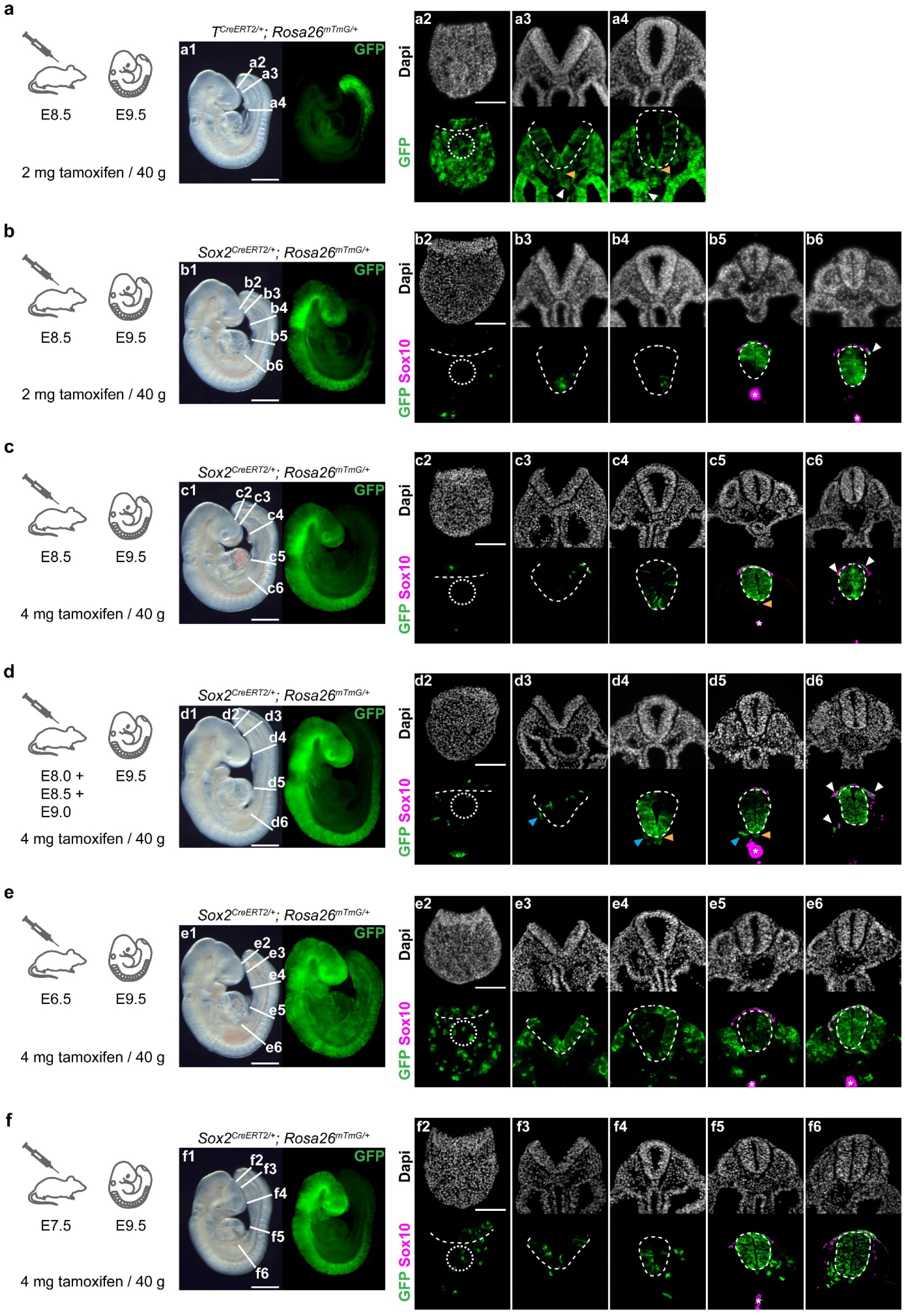
Cells expressing *Sox2* after gastrulation do not form paraxial mesoderm nor localise to the CNH. (**a**) Lineage tracing of *T*-expressing cells in embryos from *T^CreERT2/+^* x *Rosa26^mTmG/mTmG^* matings. GFP-positive cells colonise the CNH (dotted circle in **a2**) at E9.5, 24 h after tamoxifen administration, as well as neural tube and paraxial mesoderm (**a3 – 4**) in 8/8 embryos. In addition, they are consistently found in notochord and hindgut (orange and white arrowheads respectively, **a3 – 4**). (**b – d**) Lineage tracing of Sox2-expressing cells in embryos from *Sox2^CreERT2/+^* x *Rosa26^mTmG/mTmG^* matings. Sox2-expressing cells colonise the neural tube only when activated after E8.0, and do not specifically colonise the CNH, independent of tamoxifen concentration and injection time point (**b**, n = 6 embryos; **c**, n = 8 embryos; **d**, n = 12 embryos). Only single GFP-positive cells are found outside the neural tube and most co-express the neural crest marker Sox10 (white arrowheads in **b6, c6, d6**). After multiple tamoxifen injections (E8.0 – E9.0), a very low number of GFP-positive/Sox10-negative cells are found outside the neural tube (blue arrowheads in **d3 – 5**). Sox2-positive cells also colonise the notochord (orange arrowheads in **c5, d4 – d5**). (**e, f**) Tamoxifen administration to *Sox2^CreERT2/+^; Rosa26^mTmG/+^* embryos at early (E6.5) or late (E7.5) gastrulation with analysis at E9.5. GFP-positive cells colonise both neural tube and paraxial mesoderm after tamoxifen injection at E6.5 (**e**, n = 12/12 embryos), and a small number of mesoderm cells are GFP-positive after tamoxifen administration at E7.5 (**f**, n = 10/10 embryos). White dotted circles in **a2 – f2** indicate the CNH. White dashed lines in **a2 – f6** outline the neuroepithelium. See Supplementary Table 2 for summary of lineage tracing experiments **b – d**. White asterisks in **b5 – 6, c5, d5, e5 – 6, f5** indicate non-specific trapping of secondary antibody in hindgut lumen. Scale bars: 500 μm for whole embryos (**a1 – f1**); 100 μm for sections (**a2 – f2**).

We next performed the reverse experiment using a *Sox2^CreERT2/+^* driver crossed with the *Rosa26^mTmG/mTmG^* reporter (Figure 4b). While the neural tube was strongly labelled in all cases, sectioning the embryo from hindbrain to tail-bud tip revealed intermittent labelling of the notochord, but very few or no GFP-positive cells in mesoderm or CNH. Those few GFP-positive cells that were located in the paraxial mesoderm were Sox10-positive indicating neural crest identity and therefore originating from the neural tube. Even when the tamoxifen dose was doubled (Figure 4c) and pregnant dams were given multiple injections of high dose tamoxifen to achieve maximal recombination (Figure 4d), a similar result was obtained: GFP-positive cells were present in neural tube and neural crest but not somites. Overall, few GFP-positive/Sox10-negative cells occurred in paraxial mesoderm (blue arrowheads in Figure 4d3 – 5), with serial sections revealing 1 – 23 GFP-positive/Sox10-negative cells per embryo (Supplementary Table 2). Hence, contribution of *Sox2*-expressing cells to paraxial mesoderm is an exceptional occurrence after E8.5.

As a positive control experiment, we administered tamoxifen prior to E8.5, at either early gastrulation (E6.5; Figure 4e) or late gastrulation (E7.5; Figure 4f). Here, GFP-positive cells contributed to both neural tube and mesoderm, especially following E6.5 tamoxifen, and to a lesser extent after E7.5 tamoxifen. This was expected as the epiblast, which forms all three germ layers during gastrulation, expresses *Sox2^26^*. We conclude that, from E8.5 onwards, *Sox2*-expressing cells are on the lineage to neural tube only.

### *Sox2* mRNA is detectable in the neural plate but not in the CNH

The hypothesis that NMPs co-express Sox2 and *T* is based partly on immunostaining in mouse^10^, chick^9^, and zebrafish embryos^11^. These studies suggest the presence of Sox2-positive/T-positive cells in the region where NMPs are believed to reside. In addition, WISH against *Sox2* and *T* on whole embryos appears to confirm this^9, 11, 20, 27^. Indeed, when we performed WISH for *Sox2* and *T* using E8.5 (Figure 5a) and E9.5 embryos (Figure 5b), the markers seemed to overlap, as described before. However, sectioning of the hybridised embryos revealed that *Sox2* is absent from the CNH (Figure 5b10), and is expressed in neuroepithelium only. On the other hand, *T* is expressed not only in the pre-somitic mesoderm and CNH, but also in the forming neural tube at both E8.5 (Figure 5a1 – 5) and E9.5 (Figure 5b1 – 5). The presence of *T* mRNA in the neural plate is surprising, yet explains why lineage tracing of *T*-expressing cells from E8.5 leads to significant contribution to the neural tube (Figure 4a3 – 4).

**Figure 5.**
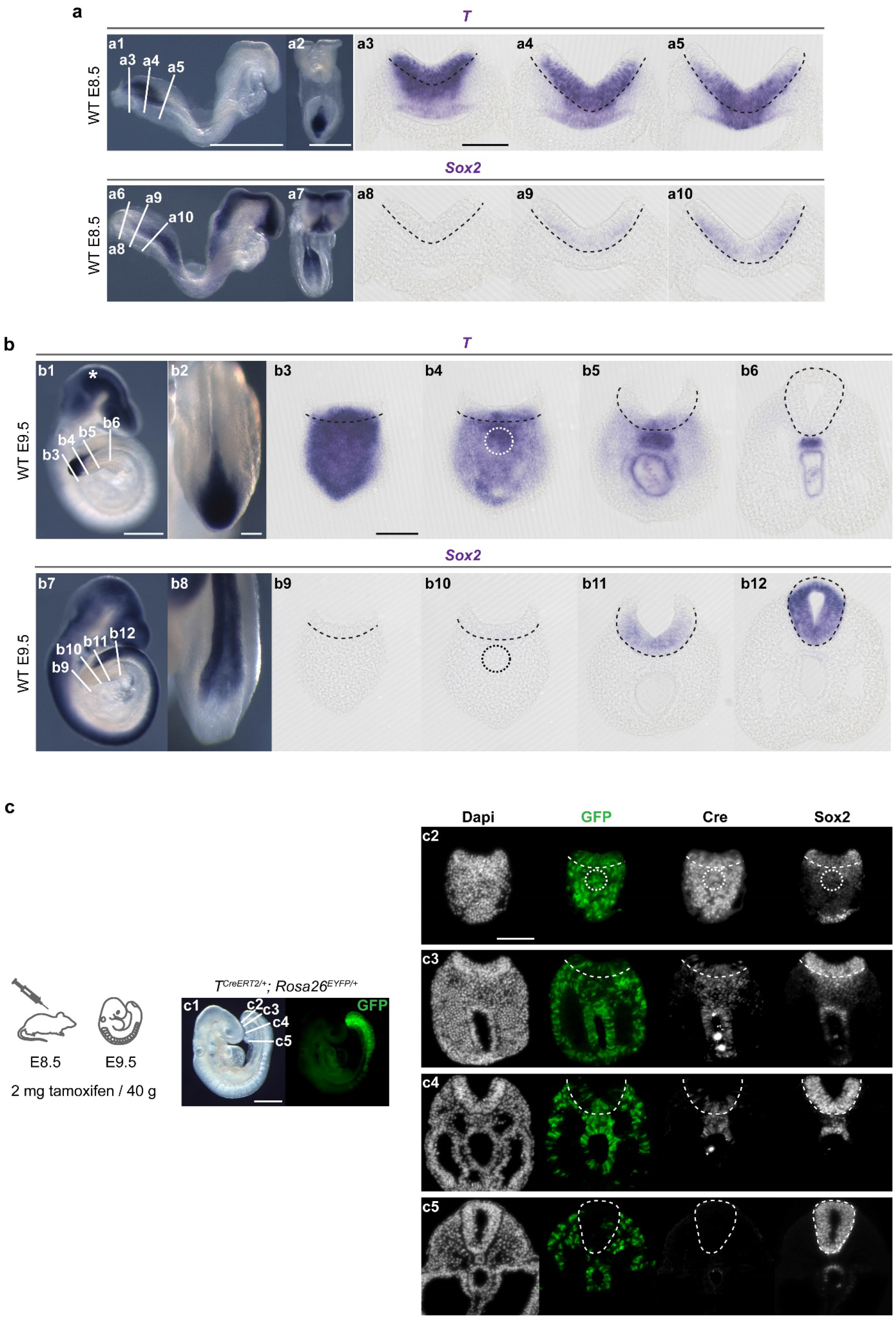
Sox2 protein but not mRNA is expressed in the CNH. (**a, b**) WISH in E8.5 (**a**) and E9.5 (**b**) wild-type (WT) embryos for *T* (n = 4, 5) and *Sox2* (n = 5, 5) mRNA. Dotted circles in **b4** and **b10** indicate the CNH. Black dashed lines in **a3 – 5, a8 – 10, b3 – 6, b9 – 12** outline the basal surface of the neuroepithelium. (**c**) Immunostaining for Cre recombinase and Sox2 in E9.5 in *T^CreERT2/+^; Rosa26^EYFP/+^* embryos (n = 7) to detect the distribution of *T* and Sox2 proteins. The pattern of Cre expression is similar to *T* mRNA expression at E9.5 (**b3 – 6**). Sox2 protein is present in the CNH (white dotted circles in **c2**) in contrast to the absence of mRNA from this location (**b10**). White asterisk in **b1** indicates non-specific trapping of the probe in the head. Scale bars: 500 μm for whole embryos (**a1 – 2, b1, c1**); 100 μm for caudal region (**b2**) and sections (**a3, b3, c2**).

T-protein expression was determined indirectly, by immunostaining for Cre recombinase in embryos from *T^CreERT2/+^* x *Rosa26^EYFP/EYFP^* matings, as no available *T* antibody gave a sufficiently specific signal. In E9.5 sections, (**T**)Cre protein expression (Figure 5c) correlated closely with the pattern for *T* mRNA (Figure 5b3 – 6), lending support to this indirect T-protein methodology. (T)Cre is robustly expressed in the CNH, as is Sox2, albeit with lower intensity, in correspondence with previous reports^9, 11, 20^. Presumably, high Sox2 protein stability, which is estimated at > 48 h^28–32^, is responsible for this residual Sox2 immunostaining, that likely stems from gene expression at the epiblast stage.

### Deleting *Sox2* in the *T*-expressing lineage does not affect mesoderm formation

If *Sox2* expression is limited to cells with neuroepithelial fate, from E8.5 onwards, then mesoderm should be unaffected by deletion of *Sox2* specifically in the *T*-expressing lineage. To test this prediction, *T^CreERT2/+^*; *Sox2^fl/+^* x *Sox2^fl/fl^* matings were performed and CreERT2 was activated at E7.5, with embryo collection at E9.5 (Figure 6a). There was no detectable difference in somite stage, body length or PNP length between *T^CreERT2/+^; Sox2^fl/fl^* embryos, which lack *Sox2* expression in the *T*-domain, and other genotypes within these litters (Figure 6b – d). Moreover, none of the embryos had obvious neural tube, mesodermal or other axial defects. *Sox2* expression was almost completely absent from the caudal end of *^CreERT2/+^*; *Sox2^fl/fl^* embryos (Figure 6e15 – 16, n = 4/4), whereas these embryos showed normal *Dll1* and *Pax1* expression (Figure 6f13 – 16) suggesting that mesoderm formation is indeed unaffected. Analysis of *Sox2* and *Dll1/Pax1* expression 24 h after CreERT2 activation showed a very similar phenotype, without obvious defects of neural tube or mesoderm formation (Supplementary Figure 3).

**Figure 6.**
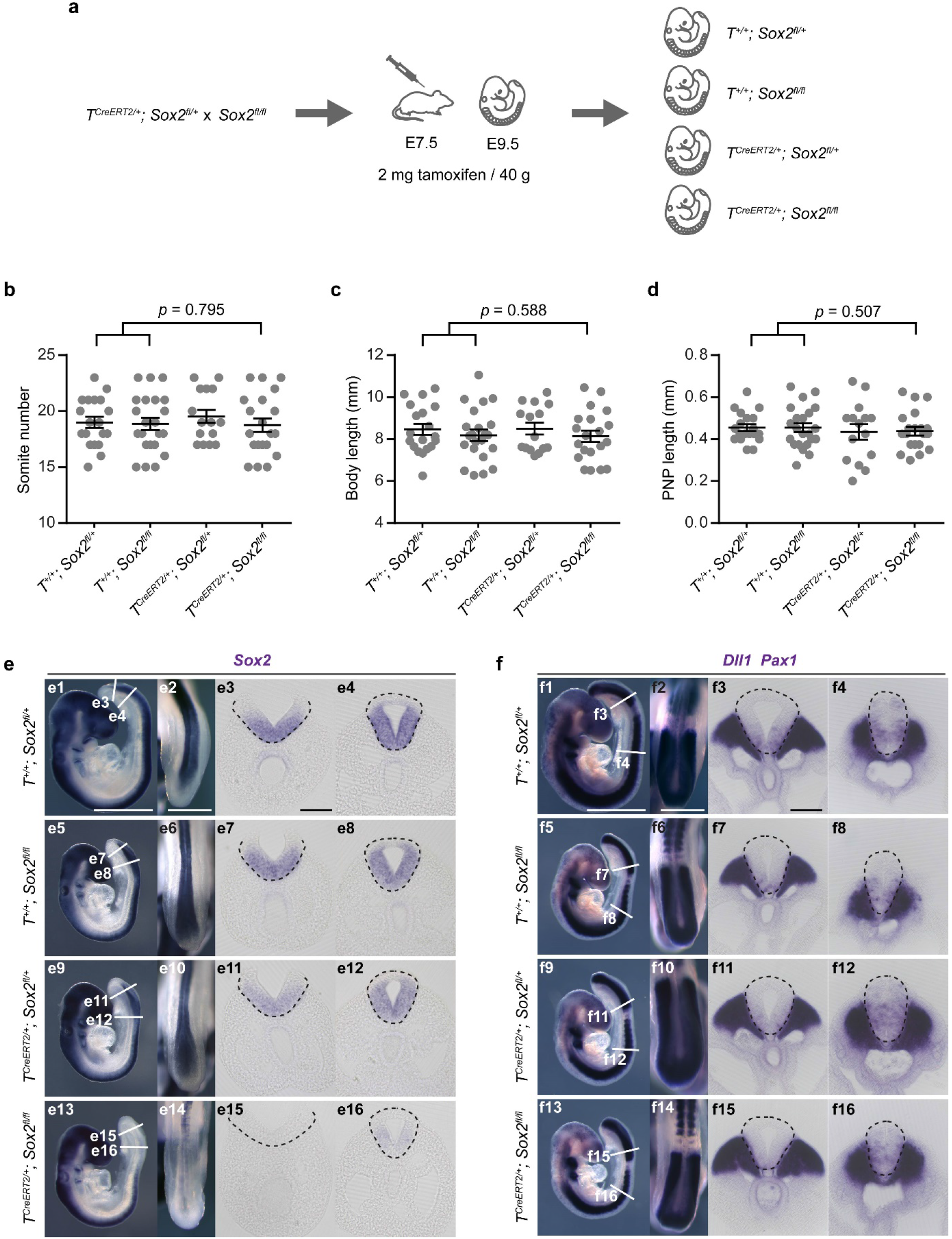
*T^CreERT2/+^*; *Sox2^fl/fl^* embryos have a *Sox2*-negative neural tube, but no mesoderm or axis elongation defects. (**a**) Specific deletion of *Sox2* in the *T*-expressing lineage in *T^CreERT2/+^*; *Sox2^fl/fl^* embryos. Tamoxifen was injected at E7.5 and embryos collected 48 h later at E9.5. Deleting *Sox2* in the *T*-expressing lineage has no effect on somite number (**b**), body length (**c**), or PNP length (**d**). Data shown as mean ± SEM including the individual data points. Unpaired, two-tailed Student’s *t*-test comparing *T^CreERT2/+^*; *Sox2^fl/fl^* embryos with CreERT2-negative embryos *(T^+/+^; Sox2^fl/+^*, n = 19; *T^+/+^; Sox2^fl/fl^*, n = 22; *T^CreERT2/+^; Sox2^fl/+^*, n = 15; *T^CreERT2/+^; Sox2^fl/fl^*, n = 20). (**e**) WISH against *Sox2* (n = 4 embryos per genotype). *Sox2* expression is almost absent from the neural tube of *T^CreERT2/+^; Sox2^fl/fl^* embryos, although neural tube morphology appears normal (**e15 – 16**). (**f**) WISH for *Dll1* and *Pax1*. Distal sections (**f3, 7, 11, 15**) show *Dll1* expression in presomitic mesoderm. Proximal sections (**f4, 8, 12, 16**) show *Pax1* expression in sclerotome. Mesoderm formation appears unaffected in *T^CreERT2/+^; Sox2^fl/fl^* embryos (n = 4 embryos per genotype for *Dll1/Pax1*). Black dashed lines in e and f outline the neuroepithelium. Scale bars: 500 μm for whole embryos and PNPs (**e1 – 2, f1 – 2**); 100 μm for sections (**e3, f3**).

Immunostaining revealed weak, patchy, residual expression of Sox2 in the neuroepithelium of *T^CreERT2/+^; Sox2^fl/fl^* embryos (Figure 7a10, 12). Sox1 and Sox3 belong to the same subgroup of SoxB1 transcription factors and are known to have similar sequences and overlapping expression patterns with Sox2^26, 33^, suggesting possible functional redundancy. While Sox1 was absent from the caudal region of both wild-type (WT) and *T^CreERT2/+^; Sox2^fl/fl^* embryos (Figure 7a4, 6, 10, 12), Sox3 was found to be expressed throughout the WT neural tube, like Sox2, and exhibited possible up-regulation in the *T^CreERT2/+^*; *Sox2^fl/fl^* neural tube (n = 5/5). Hence, Sox3 may potentially compensate for the loss of Sox2, in ensuring normal neural tube development.

**Figure 7.**
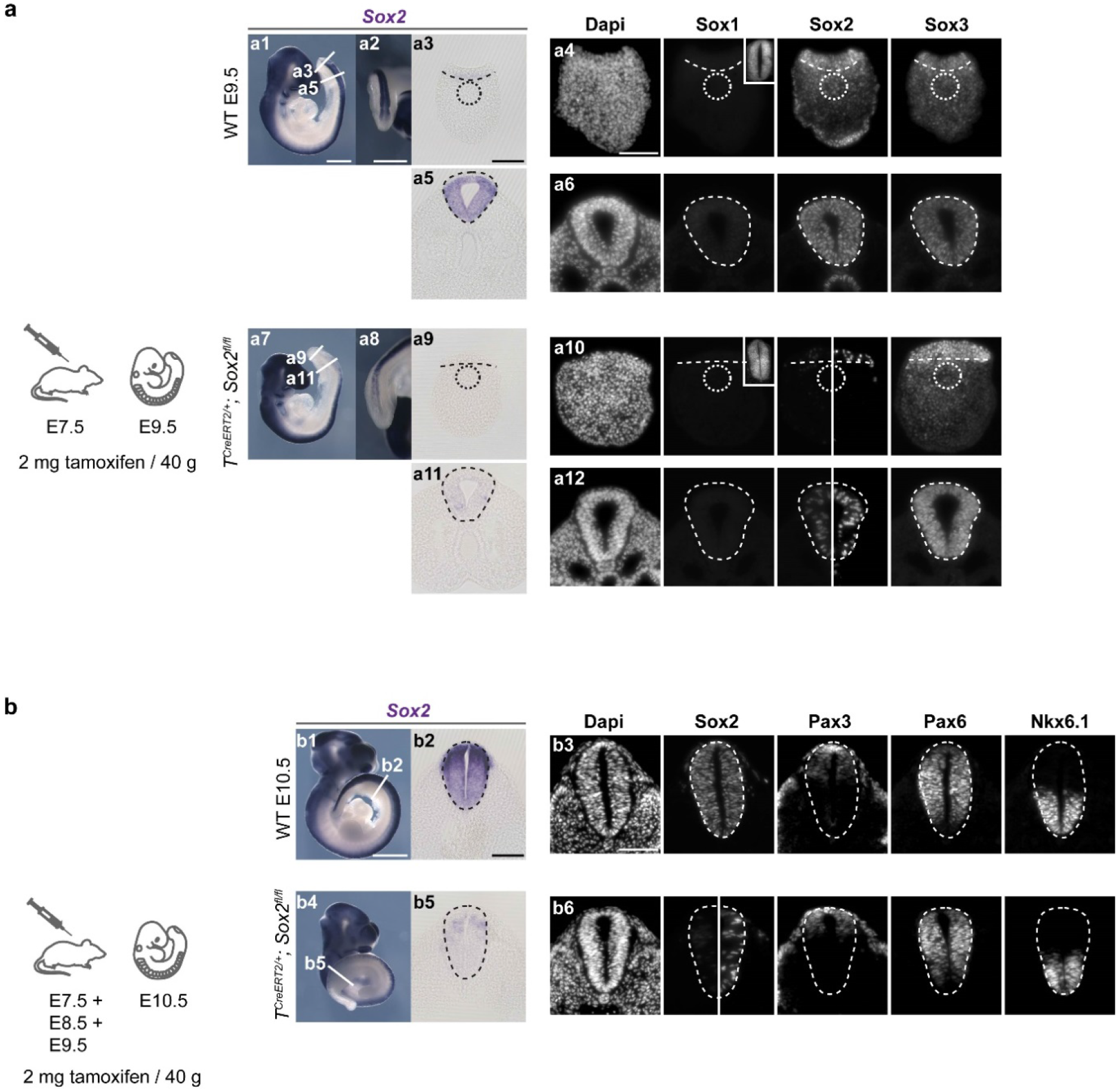
*T^CreERT2/+^; Sox2^fl/fl^* embryos form a normal neural tube. (**a**) Sox2 protein and mRNA levels are down-regulated in n = 5/5 (immunostaining) and n = 4/4 (WISH) *T^CreERT2/+^; Sox2^f^* embryos. Sox2 staining: left side taken with the same exposure time as for wild-type (WT); right side taken with longer exposure time. Sox1 is not expressed at the caudal end of either WT nor in *T^CreERT2/+^; Sox2^fl/fl^*embryos. Insets in **a4** and **a10** show cranial sections from the same embryo, stained and processed in parallel, as positive control for Sox1 antibody. Sox3 appears up-regulated in n = 5/5 *T^CreERT2/+^*; *Sox2^fl/fl^* embryos compared to WT (n-values for WISH, IHC: WT, n = 3, 5 embryos; *T^CreERT2/+^; Sox2^fl/fl^*, n = 4, 5 embryos). Dotted circles in **a3 – 4** and **a9 – 10** indicate the CNH. (**b**) Following multiple tamoxifen injections, n = 4/4 *T^CreERT2^+; Sox2^flfl^* embryos show normal neural tube morphology with normal dorso-ventral expression domains of Pax3, Pax6 and Nkx6.1 (n-values for WISH, IHC: WT, n = 3, 3 embryos; *T^CreERT2/+^; Sox2^fl/fl^*, n = 3, 4 embryos). Dashed lines in **a3 – 6, a9 – 12, b2 – 3, b5 – 6** outline the neuroepithelium. Scale bars: 1 mm for whole embryo (**b1**), 500 μm for whole embryo and PNP (**a1 – 2**); 100 μm for sections (**a3 – 4, b2 – 3**).

We attempted to further deplete *Sox2* expression in the *T*-expressing lineage, by giving pregnant dams multiple tamoxifen injections between E7.5 and E9.5, with embryo collection at E10.5 (Figure 7b). Compared with WT controls, *T^CreERT2/+^; Sox2^fl/fl^* embryos did not display any visible morphological abnormalities (n = 0/15). Although *Sox2* mRNA expression was clearly reduced along the caudal part of these embryo, it was still detectable in sections, albeit with markedly reduced intensity (Figure 7b4 – 5; n = 3/3). Similarly, Sox2 protein expression was further reduced but still detectable after long exposure times (Figure 7b6; n = 4/4). Hence, it was not possible to entirely eliminate *Sox2* expression, by recombination in the *T*-expression domain, suggesting that the caudal neural tube may be derived in part from *T*-negative cells. Immunostaining for Pax3, Pax6 and Nkx6.1, as markers of dorsal, lateral and ventral neural tube cell populations revealed that initial neural tube specification is unaffected by elimination of most *Sox2* expression (Figure 7b6; n = 4/4 embryos).

### Deleting *Sox2* in the *T*-expressing lineage causes NMPs to accumulate in the tail-bud

Finally, we investigated NMP behaviour in *T^CreERT2/+^; Sox2^fl/fl^* embryos. CreERT2 was activated at E7.5, with embryo collection at E8.5, followed by DiO injection into region 1 of the CLE. After 24h culture to E9.5, DiO-positive cells were detected in the CNH of both *T^CreERT2/+^; Sox2^fl/fl^* and *T^+/+^*; *Sox2^fl/+^* control embryos (white arrowheads in Figure 8a2, 8a3, 8a5), with contribution to both paraxial mesoderm and dorsal neural tube (Figure 8a6). Strikingly however, 8/9 *T^CreERT2/+^*; *Sox2^fl/fl^* embryos exhibited an apparent accumulation of DiO-positive cells at the caudal-most end of the body axis (dotted magenta line in Figure 8a3; 8a4). This was not seen in control embryos without CreERT2 expression (dotted grey line in Figure 8a2; n = 0/6). This finding suggests that some *Sox2*-depleted cells may not be able to adopt a neural fate and instead accumulate in the caudal-most axial extremity, where exclusively mesodermal progenitors are located.

**Figure 8.**
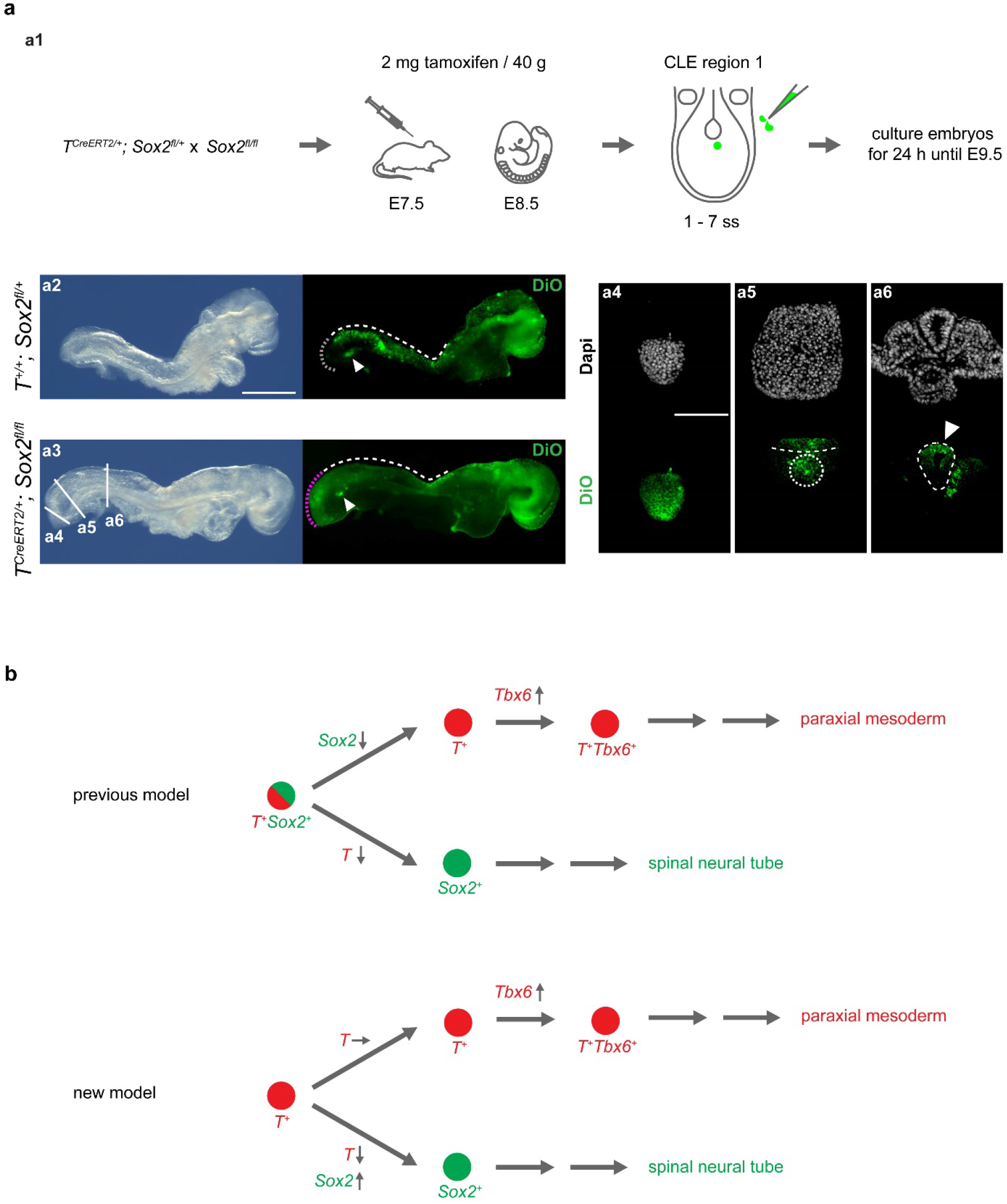
DiO-labelling shows NMP cell accumulation in the tail-bud of *T^CreERT2/+^; Sox2^fl/fl^* embryos. (**a**) DiO-labelling of CLE region 1 in embryos from *T^CreERT2/+^*; *Sox2^fl/+^* x *Sox2^fl/fl^* litters, 24 h after tamoxifen administration at E7.5. DiO-positive cells colonise the CNH of both *T^+/+^; Sox2^fl/+^* and *T^CreERT2/+^; Sox2^fl/fl^* embryos (white arrowheads in **a2, a3**; n = 6/6 and 8/9). Sections show DiO-positive cells in dorsal neural tube of *T^CreERT2/+^; Sox2^fl/fl^* embryos (white arrowhead in **a6**, n = 9/9) and in paraxial mesoderm. DiO-labelled cells also form a ventral extension of the caudal-most tail-bud of *T^CreERT2/+^; Sox2^fl/fl^* embryos (dotted magent line in **a3**; shown in section in **a4**; n = 8/9), a location where exclusively mesodermal progenitors reside. This is not seen in *T^+/+^; Sox2^fl/+^* control embryos (dotted grey line in **a2**; n = 0/6). E9.5 embryos in this experiment had normal somite numbers but a relatively short body axis, likely due to combined stress of tamoxifen and embryo culture. White dashed lines in **a2 – 3** indicate rostral extent of DiO-positive cell colonisation. White dashed lines in **a5 – 6** outline neuroepithelium. (**b**) Previous and revised NMP lineage models. See text for explanation. Scale bars: 500 μm for whole embryo (**a2**); 100 μm for sections (**a4**).

## DISCUSSION

In 1884, Kölliker postulated that the spinal neural tube is mesoderm-derived^34^. Yet, this idea fell into oblivion until Cambray and Wilson performed their grafting studies and identified two regions in the early mouse embryo that give rise to both neural tube and somites^1, 2^, suggesting the possibility of a shared progenitor. Our study refines their fate map and confirms that the E8.5 caudal region harbours two closely adjacent populations with NMP-like characteristics, the NSB and CLE region 1. Both areas fulfil the criteria of NMPs, in that their progeny are located in the CNH at E9.5 as well as colonising both neural tube and somites along extended axial lengths. Yet, labelling of CLE region 1 at E8.5 reveals colonisation of paraxial mesoderm and dorsal neural tube, whereas the NSB, which lies directly rostral to CLE region 1, contains cells that contribute to paraxial mesoderm, notochord and ventral neural tube. In line with these findings, Cambray and Wilson consistently found contribution to dorsal neural tube and somites, but sometimes also to the notochord when transplanting the rostral CLE^2^, suggesting their grafts may have included cells from both NSB and CLE region 1.

This highlights the need for clear terminology, although various terms are used inconsistently in different studies of NMPs. For example, some authors use the terms NSB and CLE interchangeably when referring to the NMP location^10, 13, 20^, although these regions contain populations with different colonisation patterns. In other studies, the authors refer to the entire CLE as the source of NMPs^10, 19, 35–38^, and not just the rostral aspect. A similar issue applies to the CNH, where NMPs are located after E9.0. When viewing transverse embryonic sections in a rostral-to-caudal direction we found that, after labelling CLE region 1, DiO-positive cells were specifically retained in the first ~3 sections beyond the level where the hindgut lumen is lost. This is the likely location of the CNH, and yet several authors show sections they consider to contain the CNH which are clearly rostral to this level, as the notochord and hindgut lumen are visible^18, 20, 22, 36^.

Strikingly, *in vivo* lineage tracing reveals that Sox2-positive cells do not in fact contribute to paraxial mesoderm after gastrulation. Given that NMPs are proposed to contribute to mesodermal and neural lineages over several days of development, as the body axis elongates^22, 39^, it seems unlikely that *T/Sox2* double-positivity is an inherent property of NMPs. Instead, our data suggest that NMPs are contained only in the T-expressing cell population. Moreover, axis elongation is unaffected when *Sox2* is deleted in the *T* lineage. Taken together, these findings suggest a model in which *T* is expressed in cells of both mesodermal and neural fate, whereas *Sox2* functions downstream of T, solely in the neural lineage (Figure 8b). This model receives support from analyses of mutant mice with axial defects and premature axis truncation^22^. In some (e.g. *Wnt3a* and *Tbx6* mutants), ectopic neural tubes are formed instead of mesoderm^40–45^ from the level of the sixth somite onwards, i.e. the axial level from which NMPs are proposed to give rise to somites and neural tube^39, 46^. However, the converse phenotype, of ectopic paraxial mesoderm in the absence of neural tube formation, does not appear to have been described, supporting the idea that neural fate is ‘downstream’ of mesodermal fate in NMP differentiation. This is further underpinned by our laser ablation experiments, which showed that ablation of the NMP region resulted in embryos with transient excess disorganised neural tissue, but strongly decreased expression of pre-somitic mesodermal markers. On the other hand, cells traced from the NMP location accumulated in the tail-bud tip of embryos lacking *Sox2* in the *T*-expression domain. As this region exclusively contains mesodermal progenitors, these findings suggest that some NMPs are directed towards a mesodermal fate in the absence of *Sox2*.

As well as producing no discernible defects of paraxial mesoderm formation or axial elongation, ablation of *Sox2* in *T*-expressing cells had no apparent effect on neural tube morphology or dorso-ventral patterning. This is a striking result, given the severe effects of complete *Sox2* loss during development^28^. It is possible that other SoxB1 family members compensate for lack of Sox2^47, 48^, and indeed we observed up-regulation of Sox3 in *Sox2-* depleted embryos, similar to a previous study^49^. Moreover, even the most stringent tamoxifen regimes were unable to completely remove *Sox2* expression when recombined in the *T*-expression domain, suggesting that *T*-negative cells may also contribute to the neural tube, albeit in small numbers.

When lineage tracing *T*-expressing cells at E8.5, we detected a contribution to the hindgut, a structure that is usually assumed to be derived from endoderm established at gastrulation. This was briefly mentioned previously^25^, but does not appear to have been further investigated. Moreover, lineage tracing experiments reveal that both Sox2-positive and T-positive cells contribute to the notochord, a structure that is often considered to be mesodermal^50^. These findings suggest that the hindgut may be derived, at least in part, from *T*-expressing cells, while the notochord may receive a contribution from *Sox2*-expressing cells that have embarked upon the neural lineage.

In conclusion, we have further defined the post-gastrulation embryonic domain in mouse embryos that contains cells with NMP-like behaviour, and demonstrated that these cells express T, but not *Sox2*. Deletion of *Sox2* in *T*-expressing cells does not compromise paraxial mesoderm formation or axial elongation, and there are only subtle neural defects, suggesting likely functional compensation by other *Sox* genes. These findings give a revised perspective on mouse NMPs that departs from the *T/Sox2* double-positive model that is currently prevalent.

## ONLINE METHODS

### Mouse procedures

Animal studies were in accordance with the UK Animals (Scientific Procedures) Act 1986 and the Medical Research Council’s ‘Responsibility in the Use of Animals for Medical Research’ (July 1993). Non-mutant embryos were from random-bred CD-1 mice. CreERT2 lines were: *T^CreERT2/+^* (Tg(T-cre/ERT2)1Lwd, stock no. 025520, Jackson Laboratory)^25^ and *Sox2^CreERT2/+^* ^51^. Floxed lines were *Sox2^fl/fl^ (Sox2^tm1.1Lan^/J;* stock no. 013093, Jackson Laboratory)^52^, the *Rosa26^mTmG/mTmG^* reporter^53^, and the *Rosa26^EYFP/EYFP^* reporter^54^. *Rosa26^mTmG/mTmG^* was used for all lineage tracing experiments in Figure 4, and *Rosa26^EYFP/EYFP^* was used for experiments in Figures 5c and 8a. Sections from both reporter lines were immunostained for GFP to enhance signal. Mice were mated overnight and checked for a copulation plug next morning, which was designated embryonic day 0.5. CreERT2 recombinase was activated by intra-peritoneal injection of tamoxifen (Sigma) into the pregnant dam. See figures for tamoxifen concentrations and injection time-points.

### Embryo collection, culture and DiO-labelling

Embryos between E8.5 and E10.5 were dissected in Dulbecco’s Modified Eagle’s Medium (Invitrogen) containing 10% fetal bovine serum (Sigma). Whole embryo culture was as described^55, 56^, with yolk sac DNA used for genotyping. The lipophilic dye DiO (Vybrant^®^ DiO Cell-Labelling Solution, Molecular Probes) was used to track cells from various locations during neural tube closure. The dye was injected into embryos using glass microinjection needles controlled by a mouth pipette. Whole embryos were imaged using a Leica MZ FLIII stereoscope with Leica DC500 camera. PNP length was measured using an eyepiece graticule on a Zeiss SV11 stereomicroscope.

### Laser ablation of the rostral CLE

Region 1 of the CLE was ablated using two-photon microscopy (ZEN 2.1 Imaging Software, black edition, Zeiss). Region 1 of the CLE was labelled with DiO in embryos with 1 – 7 somites, to identify the region of interest (ROI) under the microscope. Ablation was performed using the Zeiss LSM 880 equipped with an incubation chamber, which was set at 37 °C. The laser source was a Spectra-Physics^®^ Mai Tai^®^ eHP DeepSee™. The labelled embryo was transferred to a small petri dish with a thin layer of 1% low-melting point agarose (Sigma, #A9414) in 1x PBS and filled with dissection medium (DMEM + 10% FBS). The petri dish was placed on the microscope stage to position the embryo using the A-Plan 2.5x/0.06 objective (Zeiss). Ablation was performed in a single z plane in an ROI encompassing the DiO labelled cells, with a W Plan-Apochromat 10x/0.50 objective (Zeiss) at 800 nm with 100% laser power and maximum scan speed (1.66 μm pixels, pixel dwell of 0. 77 μs) using 100 iterations (approximately 800 μs per iteration for a typical 60 μm diameter ROI). This ensured specific ablation and reduced any heat-induced damage. Successful ablation led to cavitation at the ROI and completely removed dye-labelled cells, which was confirmed by scanning through the z-axis of the embryo recording both DiO fluorescence and transmitted light. Ablated embryos were placed in culture, using as controls embryos that had have also been injected with DiO but did not undergo laser ablation.

### Immunofluorescence

Embryos were fixed in 4% paraformaldehyde in PBS for 2 h at 4 °C and processed for cryosectioning. Immunostaining was performed on transverse sections of 10 μm thickness. Primary antibodies: rabbit monoclonal anti-Sox2 (ab92494, Abcam, 1:500), chicken polyclonal anti-GFP (ab13970, Abcam, 1:500), mouse monoclonal anti-Sox10 (sc-365692, Santa Cruz, 1:500), mouse monoclonal anti-Pax3 (MAB2457, R&D, 1:200), mouse monoclonal anti-Cre recombinase (MAB3120, Merck Millipore, 1:1,000), rabbit monoclonal anti-Sox1 (GTX62974, Insight Biotechnology, 1:200), rabbit polyclonal anti-Sox3 (ab183606, Abcam, 1:200), rabbit polyclonal anti-Pax6 (901301, BioLegend, 1:200), mouse monoclonal anti-Nkx6.1 (F55A10, Developmental Study Hybridoma Bank, deposited by Madsen, O.D., 1:5). All secondary antibodies were purchased from Thermo Fisher Scientific and used at 1:500 dilution: goat anti-chicken IgY Alexa Fluor 488 (A-11039), goat anti-mouse IgG Alexa Fluor 488 (A-11029), goat anti-mouse IgG2a Alexa Fluor 488 (A-21131), goat anti-rabbit IgG Alexa Fluor 488 (A-11070), goat anti-mouse IgG Alexa Fluor 568 (A-11019), goat antimouse IgG1 Alexa Fluor 568 (A-21124), goat anti-rabbit IgG Alexa Fluor 568 (A-11011), goat anti-mouse IgG Alexa Fluor 647 (A-21236), goat anti-rabbit IgG Alexa Fluor 647 (A-21244). Sections were counter-stained with DAPI (0.5 μg/ml in PBS) and imaged either with a Leica DM LB or an Olympus IX71 inverted microscope.

### Whole-mount *in situ* hybridisation

Whole-mount *in situ* hybridisation was performed using digoxigenin-labelled RNA probes as described previously^57^. Whole embryos were imaged using a Leica MZ FLIII stereoscope with Leica DC500 camera, then sectioned at 40 μm thickness using a vibratome and imaged on a Zeiss Axioplan 2 microscope with Zeiss AxioCam HRc camera. The following antisense probes were used: *Sox2^58^, T^50^, Dll1^59^*, *Pax1^60^*.

### Image analysis

Images were cropped, adjusted, and analysed using Fiji image processing software^61^. Only linear adjustments were applied equally to the image. Translocation of DiO-labelled cells after injecting CLE region 1 (Supplementary Figure 2) was analysed in transverse sections of labelled embryos which were counter-stained with DAPI (0.5 μg/ml in PBS). First, the length of the neuroepithelium was measured along its basal side between the ventral midline and the most dorsal region. This served as a template to determine the relative position of the labelled cells in that section by calculating the distance between the ventral and dorsal borders of the DiO-positive cell group and the ventral midline. To remove the influence of overall neural tube size, which varied along the axis and between embryos, the position of labelled cells was expressed as a percentage of total neuroepithelial length and then used for statistical analysis.

### Statistical analysis

Statistical analysis was carried out using IBM^®^ SPSS^®^ Statistics (version 23). The exact *p*-values are shown in the figures and details of the tests performed are given in the figure legends.

## ACKNOWLEDGEMENTS

We thank Sandra De Castro, Ana Rolo, and Paula Alexandre for helpful discussions and critical reading of the manuscript. This work was funded by a Wellcome 4-year PhD studentship (105379) and Wellcome programme grant 087525 (to A.J.C. and N.D.E.G.).

## AUTHOR CONTRIBUTIONS

A.J.C. conceived the study. A.J.C., D.M., D.S., M.A.M., J.P.M.-B., and N.D.E.G designed the experiments. D.M. performed all experimental procedures and data analyses. D.A.M. set up and optimised the laser ablation procedures and developed the macro for image analysis. D.M. and A.J.C. wrote the manuscript with contributions from all authors.

---

**Supplementary Figure 1.**
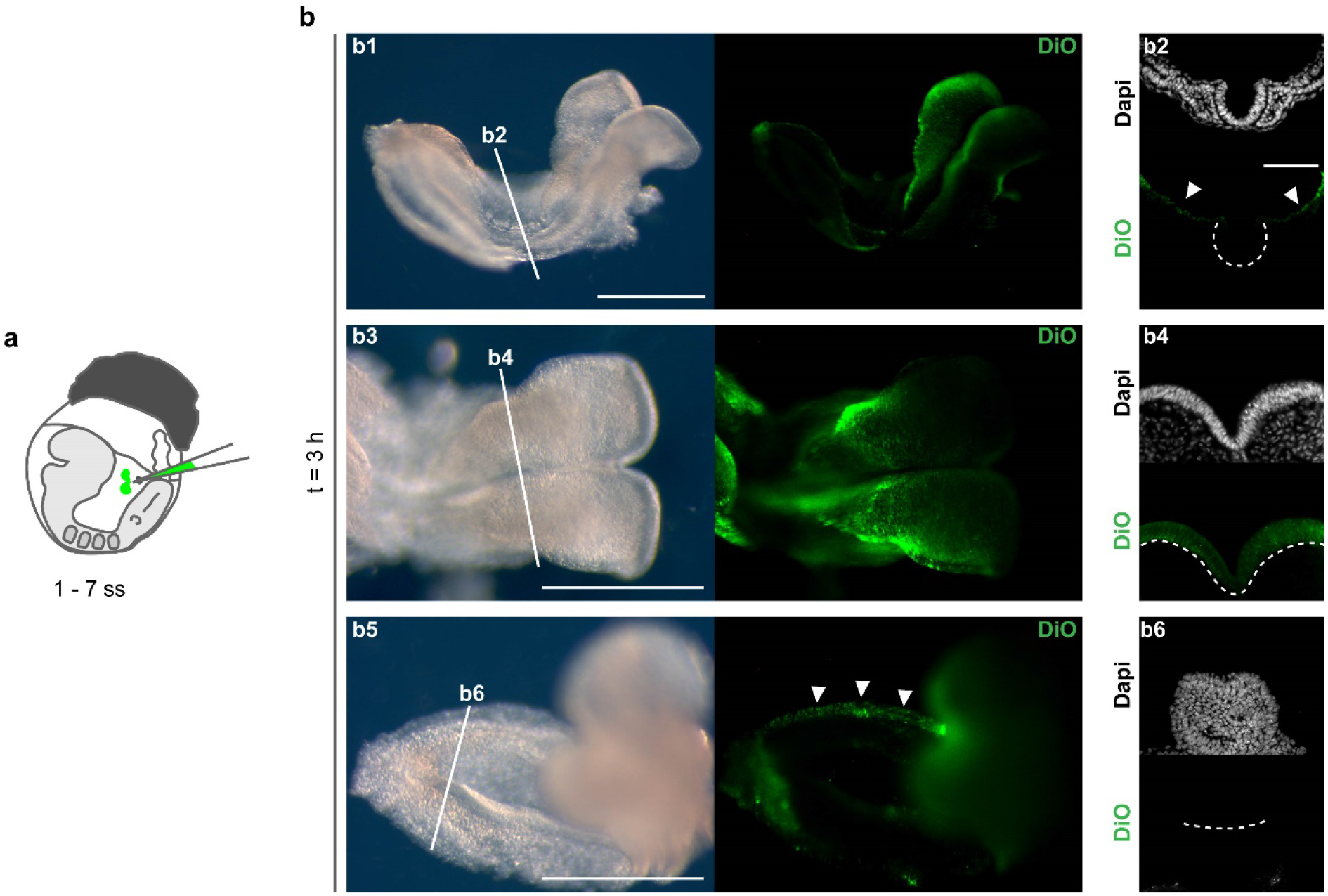
DiO binds non-specifically to embryonic head-folds. (**a**) A small amount of DiO was injected into the amniotic cavity of E8.5 WT embryos to mimic that released during the caudal labelling procedure. (**b**) After 3 h whole-embryo culture, DiO is detected extensively over the head-folds (**b3 – 4**), but not in mid-trunk (**b1 – 2**), nor tail-bud (**b5 – 6**) regions (n = 5/5). Surface ectoderm (white arrowheads in **b2**) and extraembryonic tissues (white arrowheads in **b5**) are DiO-positive in n = 5/5 embryos. White dashed lines in **b2, b4, b6** outline basal surface of the neuroepithelium. Scale bars: 500 μm for embryos (**b1, b3, b5**); 100 μm for sections (**b2**).

**Supplementary Figure 2.**
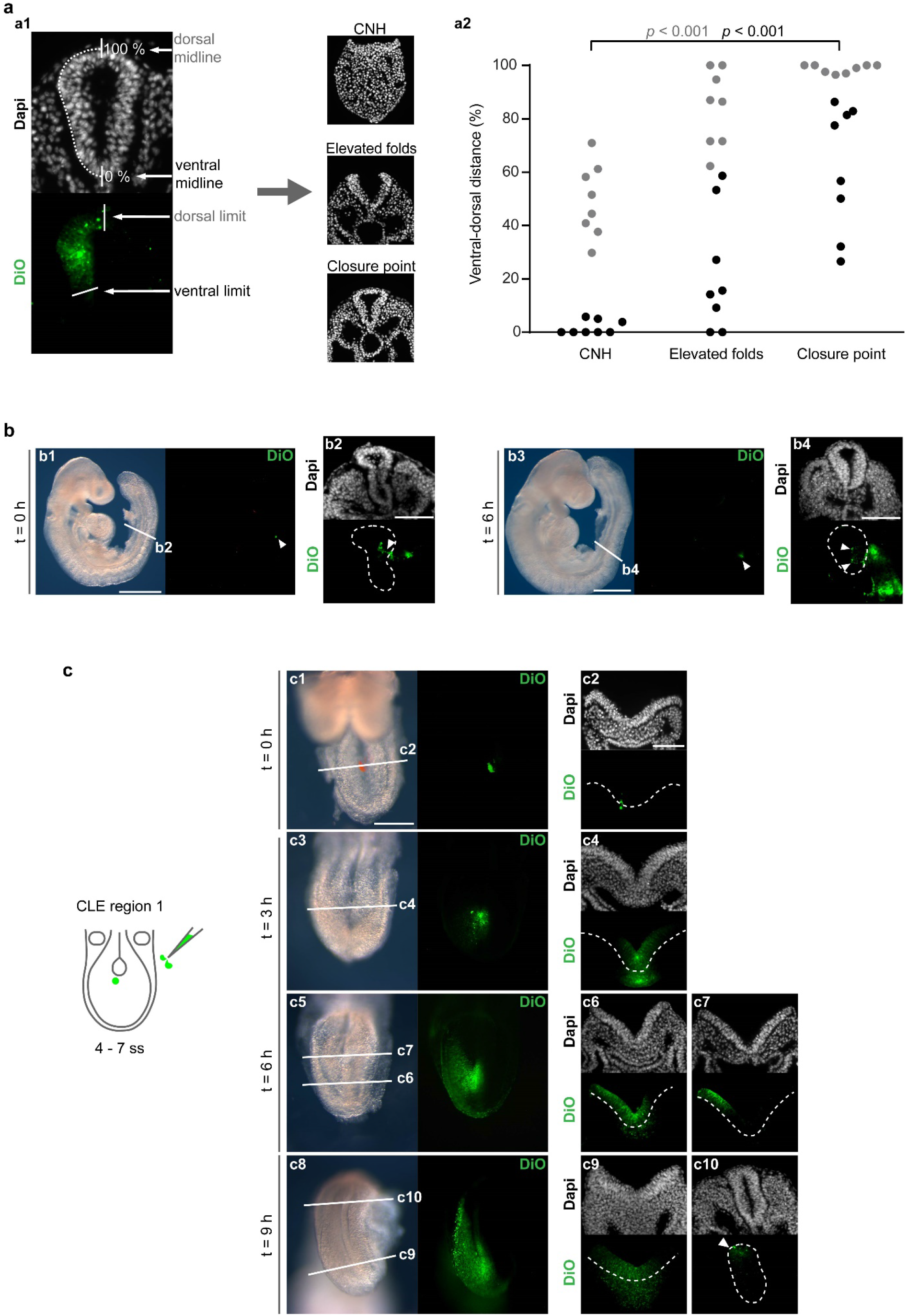
DiO-labelled cells from region 1 of the CLE specifically colonise the dorso-lateral neural tube. (**a**) Cells labelled with DiO in CLE region 1 at E8.5 translocate in a ventral-to-dorsal direction to colonise the dorso-lateral neural tube at the level of the closure point (n = 8/8 embryos). Dorsal (grey circles) and ventral (black circles) limits of DiO-positive cells are shown (**a2**) for each of 8 embryos at the three axial levels: CNH, elevated neural folds and neural tube closure point. Paired, two-tailed Student’s t-test. (**b**) Cells directly labelled with DiO in the lateral closed neural tube (white arrowheads in **b1, b2**) remain in a lateral position after 6 h whole embryo culture (white arrowhead in **b3**; in **b4** arrowheads indicate dorsal and ventral limits of DiO-labelled cells; n = 5/5 embryos). (**c**) Time-course of DiO-labelled cell translocation into the neural tube, after injecting region 1 of the CLE at E8.5 (n = 4 embryos per time point). DiO-positive cells colonise the dorsal neural tube around 9 h after injection (white arrowhead in **c10**, n = 4/4 embryos). White dashed lines in **b2, b4, c2 – 4, c6 – 7, c9 – 10** outline the neuroepithelium. Scale bars: 500 μm for embryos in (**b1, b3, c1**); 100 μm for sections (**b2, b4, c2**).

**Supplementary Figure 3.**
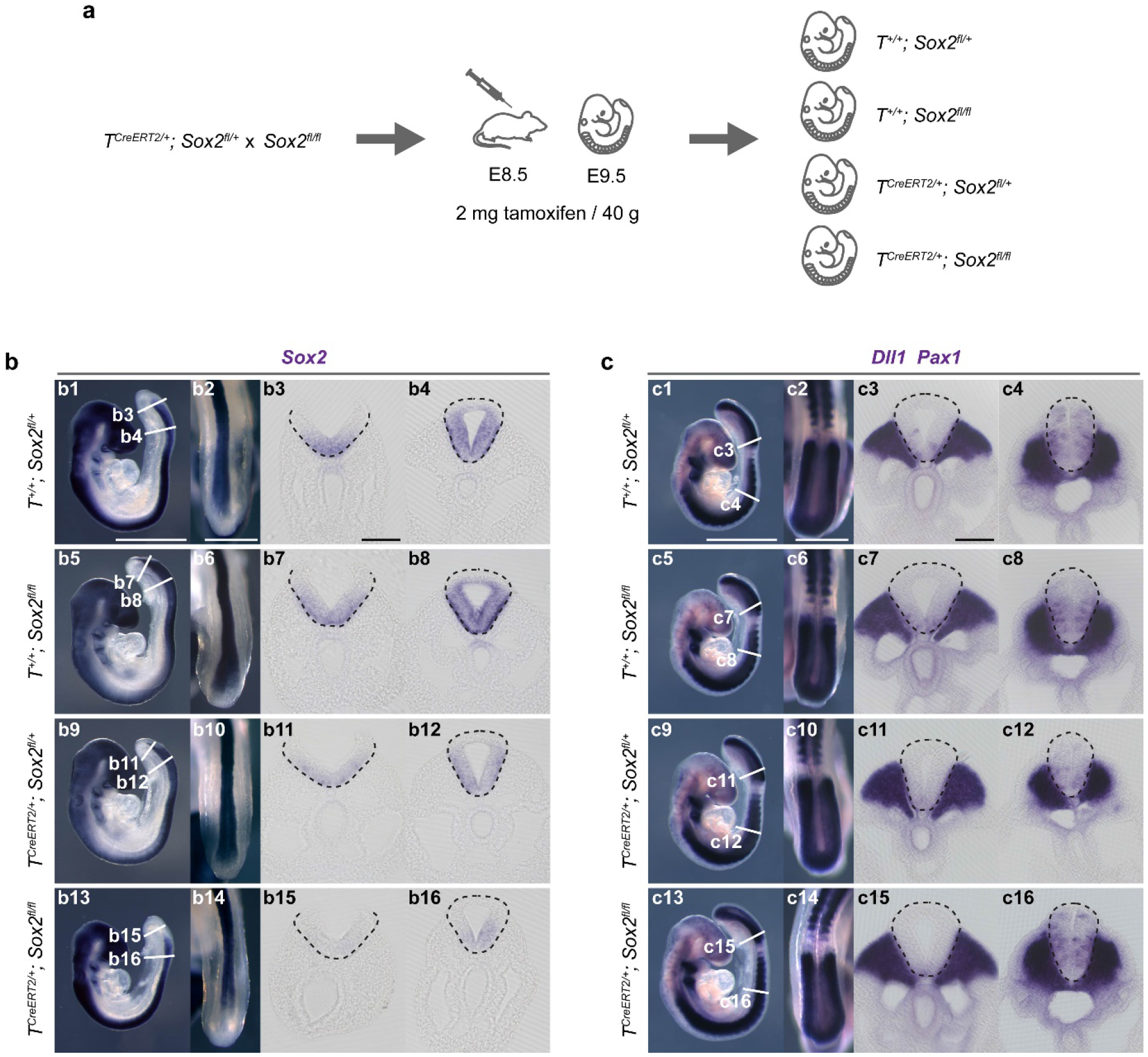
*T^CreERT2/+^*; *Sox2^fl/fl^* embryos have reduced *Sox2* expression in the caudal neural tube, but no mesoderm defects. (**a**)Specific deletion of *Sox2* in the *T*-expressing lineage, with tamoxifen injection at E8.5 and analysis at E9.5 (for comparison with Figure 6 where tamoxifen injection was at E7.5). (**b**) WISH for *Sox2* (n = 4 embryos per genotype) reveals markedly reduced *Sox2* expression in the caudal-most neural tube in *T^CreERT2/+^; Sox2^fl/fl^* embryos (**b15 – 16**; n = 4/4) compared with other genotypes. (**c**) WISH for *Dll1* and *Pax1* shows that mesoderm formation is unaffected in any of the four genotypes (n = 4 embryos each). Black dashed lines in **b** and **c** outline the neuroepithelium. Scale bars: 500 μm for whole embryos and PNPs (**b1 – 2, c1 – 2**); 100 μm for sections (**b3 – c3**).

**Supplementary Table 1.**
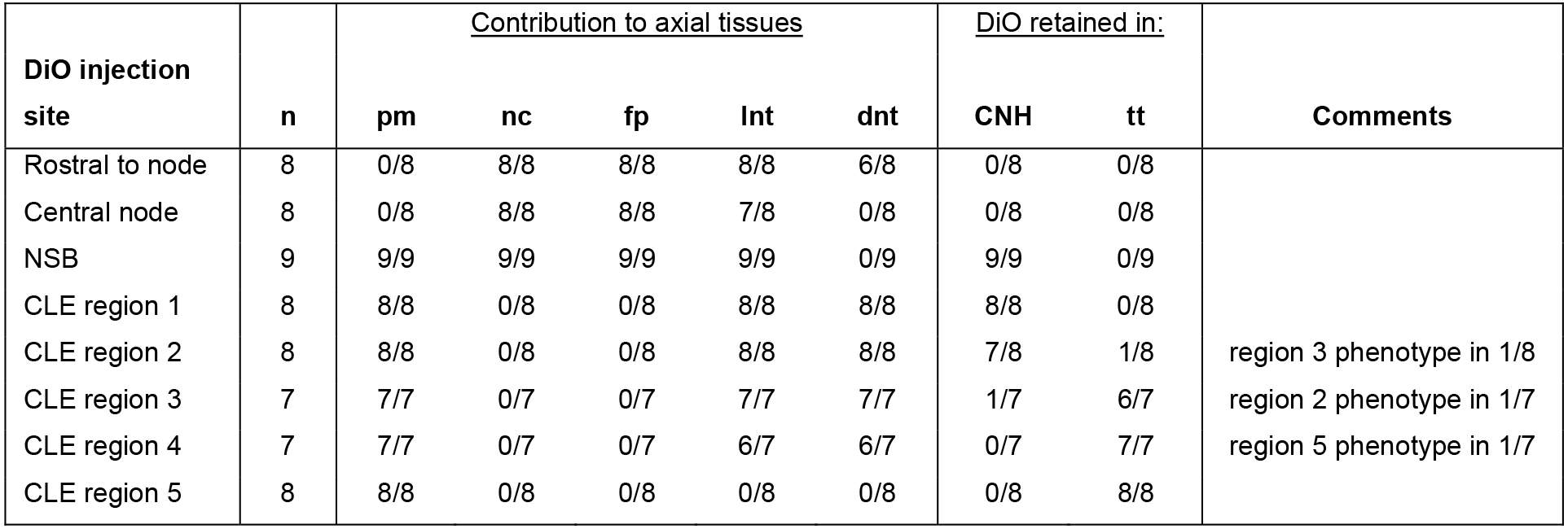
Summary of colonisation pattern of DiO-labelled cells in the experiments shown in Figures 1 and 2. Embryos were injected with DiO at E8.5 followed by 24 h whole-embryo culture. *pm*, presomitic mesoderm; *nc*, notochord, *fp*, floor-plate; *lnt*, lateral neural tube; *dnt*, dorsal neural tube; *CNH*, chordo-neural hinge; *tt*, tail-bud tip.

**Supplementary Table 2.**
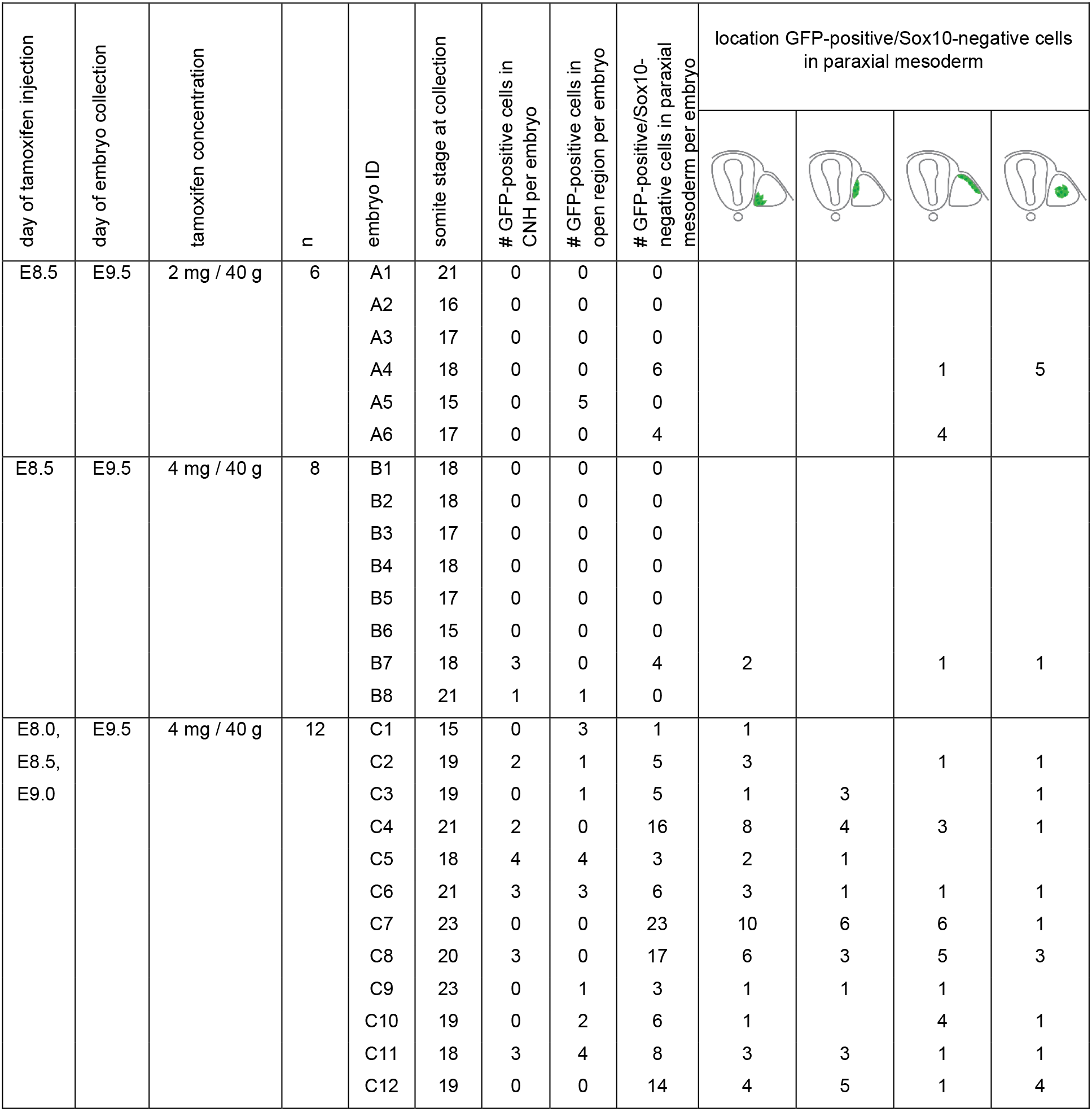
Summary of mesodermal colonisation by GFP-positive cells in *Sox2^CreERT2/+^; Rosa26^mTmG/+^* embryos, as shown in Figure 4b – d. The number of “GFP-positive cells in the open region” refers to those GFP-positive cells which were located in the presomitic mesoderm in sections at the level of the open PNP: between CNH and closure point of the neural tube. The number of “GFP-positive/Sox10-negative cells in the paraxial mesoderm” refers to the number GFP-positive/Sox10-negative (i.e. non-neural crest) cells per embryo found outside the neural tube in sections at the level of the closed neural tube.

| day of tamoxifen injection | day of embryo collection | tamoxifen concentration | n | embryo ID | somite stage at collection | # GFP-positive cells in CNH per embryo | # GFP-positive cells in open region per embryo | # GFP-positive/Sox10-negative cells in paraxial mesoderm per embryo | location GFP-positive/Sox10-negative cells in paraxial mesoderm |  |  |  |
| --- | --- | --- | --- | --- | --- | --- | --- | --- | --- | --- | --- | --- |
| E8.5 | E9.5 | 2 mg / 40 g | 6 | A1 | 21 | 0 | 0 | 0 |  |  |  |  |
|  |  |  |  | A2 | 16 | 0 | 0 | 0 |  |  |  |  |
|  |  |  |  | A3 | 17 | 0 | 0 | 0 |  |  |  |  |
|  |  |  |  | A4 | 18 | 0 | 0 | 6 |  |  | 1 | 5 |
|  |  |  |  | A5 | 15 | 0 | 5 | 0 |  |  |  |  |
|  |  |  |  | A6 | 17 | 0 | 0 | 4 |  |  | 4 |  |
| E8.5 | E9.5 | 4 mg / 40 g | 8 | B1 | 18 | 0 | 0 | 0 |  |  |  |  |
|  |  |  |  | B2 | 18 | 0 | 0 | 0 |  |  |  |  |
|  |  |  |  | B3 | 17 | 0 | 0 | 0 |  |  |  |  |
|  |  |  |  | B4 | 18 | 0 | 0 | 0 |  |  |  |  |
|  |  |  |  | B5 | 17 | 0 | 0 | 0 |  |  |  |  |
|  |  |  |  | B6 | 15 | 0 | 0 | 0 |  |  |  |  |
|  |  |  |  | B7 | 18 | 3 | 0 | 4 | 2 |  | 1 | 1 |
|  |  |  |  | B8 | 21 | 1 | 1 | 0 |  |  |  |  |
| E8.0,<br>E8.5,<br>E9.0 | E9.5 | 4 mg / 40 g | 12 | C1 | 15 | 0 | 3 | 1 | 1 |  |  |  |
|  |  |  |  | C2 | 19 | 2 | 1 | 5 | 3 |  | 1 | 1 |
|  |  |  |  | C3 | 19 | 0 | 1 | 5 | 1 | 3 |  | 1 |
|  |  |  |  | C4 | 21 | 2 | 0 | 16 | 8 | 4 | 3 | 1 |
|  |  |  |  | C5 | 18 | 4 | 4 | 3 | 2 | 1 |  |  |
|  |  |  |  | C6 | 21 | 3 | 3 | 6 | 3 | 1 | 1 | 1 |
|  |  |  |  | C7 | 23 | 0 | 0 | 23 | 10 | 6 | 6 | 1 |
|  |  |  |  | C8 | 20 | 3 | 0 | 17 | 6 | 3 | 5 | 3 |
|  |  |  |  | C9 | 23 | 0 | 1 | 3 | 1 | 1 | 1 |  |
|  |  |  |  | C10 | 19 | 0 | 2 | 6 | 1 |  | 4 | 1 |
|  |  |  |  | C11 | 18 | 3 | 4 | 8 | 3 | 3 | 1 | 1 |
|  |  |  |  | C12 | 19 | 0 | 0 | 14 | 4 | 5 | 1 | 4 |

